# Computational Evaluation of DNA Metabarcoding for Universal Diagnostics of Invasive Insect Pests

**DOI:** 10.1101/2021.03.16.435710

**Authors:** Alexander M. Piper, Noel O.I. Cogan, John Paul Cunningham, Mark J. Blacket

## Abstract

Appropriate design and selection of PCR primers plays a critical role in determining the sensitivity and specificity of a metabarcoding assay. Despite several studies applying metabarcoding to insect pest surveillance, the diagnostic performance of the short “mini-barcodes” required by high-throughput sequencing platforms has not been established across the broader taxonomic diversity of invasive insects. We address this by computationally evaluating the diagnostic sensitivity and predicted amplification bias for 68 published and novel cytochrome c oxidase subunit 1 (COI) primers on a curated database of 110,676 insect species, including 2,625 registered on global invasive species lists. We find that mini-barcodes between 125-257 bp can provide comparable resolution to the full-length barcode for both invasive insect pests and the broader Insecta, conditional upon the subregion of COI targeted and the genetic similarity threshold used to identify species. Taxa that could not be identified by any barcode lengths were phylogenetically clustered within ‘problem groups’, many arising through taxonomic inconsistencies rather than insufficient diagnostic information within the barcode itself. Substantial variation in predicted PCR bias was seen across published primers, with those including 4-5 degenerate nucleotide bases showing almost no mismatch to major insect orders. While not completely universal, a single COI mini-barcode can successfully differentiate the majority of pest and non-pest insects from their congenerics, even at the small amplicon size imposed by 2 × 150 bp sequencing. We provide a ranked summary of high-performing primers and discuss the bioinformatic steps required to curate reliable reference databases for metabarcoding studies.

## Introduction

Early detection and rapid response are crucial for preventing the establishment and spread of invasive pests and pathogens (Liebhold et al., 2016; Reaser, Burgiel, et al., 2020). Historically, invasive species surveillance has relied upon targeted inspections for predefined lists of regulated taxa (Reaser, Frey, et al., 2020; Schrader & Unger, 2003). However, as global trade networks become increasingly interlinked and anthropogenic climate change alters species range distributions, this list-based framework often lags behind the speed at which new pests can emerge and spread across borders (Bebber, 2015; Hulme, 2009). This lag becomes particularly apparent when considering impacts beyond agroecosystems, where the size and complexity of the natural environment presents challenges for accurate risk prediction (Caley et al., 2006; Crooks, 2005). In light of this, it is becoming increasingly appreciated that modern biosecurity will need to adopt a more comprehensive approach to surveillance that aims to detect and evaluate all newly introduced species, not just those regulated by national quarantine agencies (Meyerson & Reaser, 2002; Reaser et al., 2008; Simberloff, 2006). In practice, however, adoption of this framework is bottlenecked by a lack of diagnostics capacity to sort and identify the large number of specimens collected by intensive surveillance efforts (Bishop & Hutchings, 2011; Piper et al., 2019).

Plant pest and pathogen diagnostics currently rely on a mixture of morphological examination, biochemical techniques, and molecular assays such as diagnostic qPCR, and DNA barcoding (EPPO, 2019a). While these methods provide highly accurate identification for small numbers of specimens (Armstrong & Ball, 2005; Darling & Blum, 2007), their inherent restriction to analysing single specimens perreaction limits their application to large mixed samples collected in surveillance traps (Batovska et al., 2020; Carnegie & Nahrung, 2019). As an alternative, high-throughput sequencing (HTS) platforms can comprehensively characterise mixed populations of genomic DNA (metagenomics), RNA (metatranscriptomics) or taxonomically informative marker genes (metabarcoding), allowing whole communities to be identified without any prior isolation or specimen sorting (Piper et al., 2019; Tedersoo et al., 2019). While first emerging for exploring biodiversity (Handelsman, 2004; Taberlet et al., 2012), non-targeted HTS assays have recently been co-opted by various disciplines of molecular diagnostics, where their potential to act as a universal identification tool was quickly recognised (Adams et al., 2009; Comtet et al., 2015). By removing the requirement to separately develop and maintain hundreds of targeted diagnostic assays, universal HTS diagnostics could substantially expand the range of organisms within the scope of a diagnostic laboratory, as well as decrease the costs of implementation (Adams et al., 2018; Allcock et al., 2017).

Metabarcoding of the mitochondrial cytochrome c oxidase subunit 1 (COI) gene presents the most readily adoptable HTS approach for diagnostics of insect pests, due to its cost effectiveness, extensive public reference sequence databases, and ability to leverage widespread acceptance of DNA barcoding within regulatory frameworks (Andújar et al., 2018; Comtet et al., 2015; Piper et al., 2019). While conventional single-specimen DNA barcoding targets a 709 bp region of COI (Folmer et al., 1994), modern HTS platforms impose strict length limitations on sequenced molecules, and therefore “mini-barcodes” must instead be used (Brandon-Mong et al., 2015). The number of diagnostic nucleotides contained within these mini-barcodes largely determines the sensitivity and specificity for single specimens, however, mismatch between PCR primers and variable template molecules can bias amplification and cause dropouts of low-abundance taxa within mixed community samples (Elbrecht & Leese, 2015; D. W. Yu et al., 2012). Primer-template mismatch is a particular issue for protein-coding genes such as COI where variability in the third position of each codon leaves no strictly conserved regions for placement of universal PCR primers (Deagle et al., 2014). Therefore, degenerate nucleotide bases are commonly incorporated into COI metabarcoding primers to account for this inevitable mismatch (Elbrecht & Leese, 2017b), though overuse can result in undesired amplification of non-target organisms (Collins et al., 2019; Leese et al., 2021).

Historically, molecular diagnostic assays would have undergone stringent laboratory validation in order to resolve the aforementioned issues and establish performance parameters for every target designated in an unambiguously defined scope (EPPO, 2019b). However, when considering the sheer number of potential targets, hosts and matrices that would need to be evaluated for a universal assay, it is evident that many validation processes cannot be applied in their traditional sense and must be adapted to novel HTS based diagnostics (Maree et al., 2018; Roenhorst et al., 2018). In-silico methods pose a promising alternative that can leverage public reference sequence data to establish the diagnostic performance of a target barcode region and determine the best placement of degenerate PCR primers, all without requiring physical specimens (Elbrecht & Leese, 2017a; Ficetola et al., 2010). Nevertheless, the use of public reference data comes with some caveats, as issues of mislabelled taxonomic annotations, insufficiently identified specimens, and contamination are well documented (Garg et al., 2019; Locatelli et al., 2020; Pentinsaari et al., 2020; Siddall et al., 2009). It is therefore essential for public DNA barcode sequences to be appropriately curated before use for in-silico validation procedures or as reference databases for metabarcoding analysis (Piper et al., 2019).

In this article we develop a computational workflow for curating a large collection of Insect COI sequences from public sequence repositories. Using this curated database, in-silico methods are then used to evaluate the sensitivity, specificity, and predicted amplification bias for 68 published and novel metabarcoding primers on a globally relevant list of invasive insect pests and the wider insect diversity. We identify optimal subregions of the COI barcode for species differentiation and determine the amplicon lengths required to achieve comparable resolution to the full-length barcode. This study informs and offers recommendations for selection of metabarcoding primers and provides a robust workflow for assembling curated reference databases for DNA barcode-based insect identification.

## Methods

### Retrieval and curation of public reference data

COI records and mitochondrial genomes with the taxonomic annotation ‘Insecta’ were retrieved from NCBI GenBank and the Barcode of Life Data system (BOLD) (Ratnasingham & Hebert, 2007) using the *Rentrez* (Winter, David & Winter, 2017) and *bold* (Chamberlain, 2017) R packages. All retrieved sequences then went through a series of curation steps (Figure 1A). First, to resolve taxonomic synonyms between the two repositories, sequence annotations were mapped into the Open Tree of life Taxonomy (OTT) (Hinchliff et al., 2015), and only those with complete binomial names and not flagged with uncertain taxonomic placement were retained (see supplementary notes 2 and 3 for relevant flags). All sequences were then aligned to a reference Profile Hidden Markov Model (PHMM) (Eddy, 1998) of the COI locus generated from a manually curated version of the Midori-longest V237 dataset (Machida et al., 2017) using the *aphid* R package (Wilkinson, 2019). All sequences that met a minimum odds-ratio alignment score of 100 without containing stop-codons or frameshift mutations were retained, and bases outside the bounds of the LCO1490-HCO2198 (Folmer et al., 1994) primer binding sites trimmed from the alignment. To identify putatively misannotated sequences, the alignment was hierarchically clustered using the *kmer* R package (Wilkinson, 2018) and sequences removed if their species annotation at 99% identity, genus annotation at 97% identity, or family annotation at 95% identity disagreed with more than 80% of other sequences within its respective cluster. A nucleotide BLAST search (Altschul et al., 1990) was then conducted against a local contaminants database and sequences with percentage coverage and identity >79% for Wolbachia, >98% for known pseudogene sequences, or >96% for human mitochondria were removed. To identify invasive insect species within the curated sequence database, taxonomic names were retrieved from 11 global and geographically focused pest and invasive species lists (Supplementary Note 1), hereafter referred to as the ‘pest’ dataset. All sequences <200 bp were removed, then the number of barcodes per pest taxon was compared to the remainder of the insect taxa using the Welch t-test. Finally, in order to accelerate downstream computations and minimize the effects of taxonomic sampling bias (Mutanen et al., 2016), the database was pruned to a maximum 5 representative sequences per species, discarding sequences sequentially from smallest to largest.

**Figure 1:**
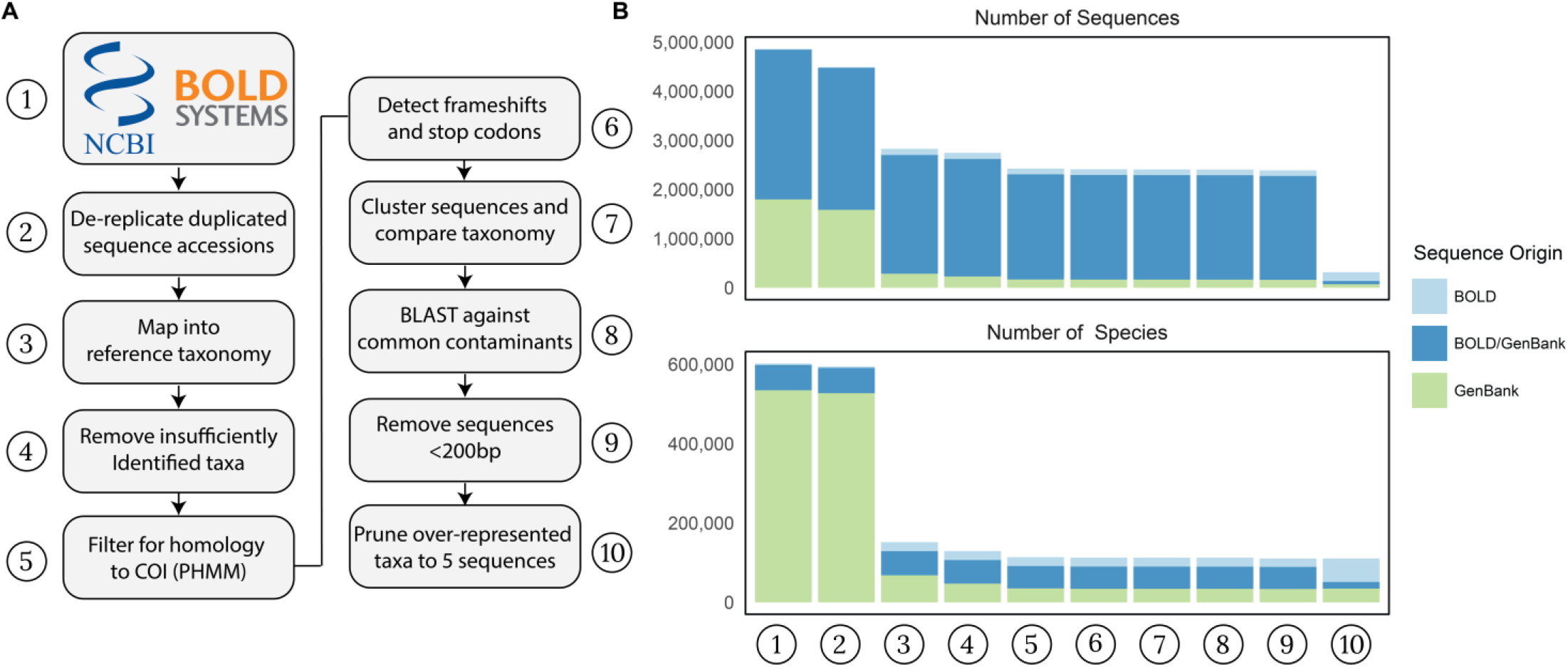
**A)** Overview of the computational pipeline used to curate public reference sequence data for primer evaluation. **B)** Number of sequences (upper) and species (lower) retained after each curation step, and their origin from BOLD, GenBank or duplicated across both repositories.

### Construction of phylogenetic tree

The curated Insect reference sequences were supplemented with an outgroup of 15 COI sequences from Diplura, the sister taxa to Insecta (Misof et al., 2014), and all positions in the alignment that contained gaps in >95% of sequences were masked. A maximum likelihood phylogenetic tree was generated using FastTree (Price et al., 2009) following the General Time-Reversible (GTR) model (Tavaré, 1986) and gamma model of rate heterogeneity across sites, with taxonomic identities at the domain, phylum, class and order ranks used to constrain the deeper topology of the tree. The constructed phylogeny was rooted on the edge connecting the Diplura outgroup to the rest of the tree, and made ultrametric using the *geiger* R package (Pennell et al., 2014) and *PATHd8* (Britton et al., 2007). All phylogenetic trees were plotted using the *ggtree* (G. Yu et al., 2017, 2018) and *ggplot2* R packages (Wickham, 2016).

### Diagnostic information within the COI barcode

The curated reference sequence database was split by taxonomic order, and Shannon’s entropy (Scheider & Stephens, 1990) calculated for each alignment position, then visualised with structural motifs annotated as per Pentinsaari et al. (2016). To identify the most diagnostically informative subregions of the COI barcode, a sliding window approach was used to split the alignment into virtual amplicon sequences of 200 bp, 300 bp and 400 bp, in intervals of 3 bp. The sequences within each virtual amplicon window were clustered at 97% similarity using *UCLUST* (Edgar, 2010), and a species considered successfully identified if there were no other sequences with conflicting taxonomic annotations within its respective cluster.

### Characteristics of published and novel primers

32 forward and 31 reverse primers (Figure 2B) overlapping the COI barcode region were identified from a literature search for the terms ‘metabarcoding mini-barcode’, and ‘metabarcoding primer’ and supplemented with 2 new forward and 3 new reverse primers designed using Primer3 (Untergasser et al., 2012). The number of reference sequences for which each primer had an appropriate binding site was determined via string matching using the *Biostrings* R package (Pagès et al., 2019), allowing for a hamming distance of 2. The frequency of each nucleotide base, as well as presence of homopolymers or GC clamps (2 or more contiguous G or C bases at the 5’ end) were determined using the *Biostrings* and *DECIPHER* R packages (Wright, 2016). Primer melting temperatures were calculated using nearest neighbour thermodynamics (SantaLucia & Hicks, 2004) with the *TmCalculator* package (Li, 2019).

**Figure 2:**
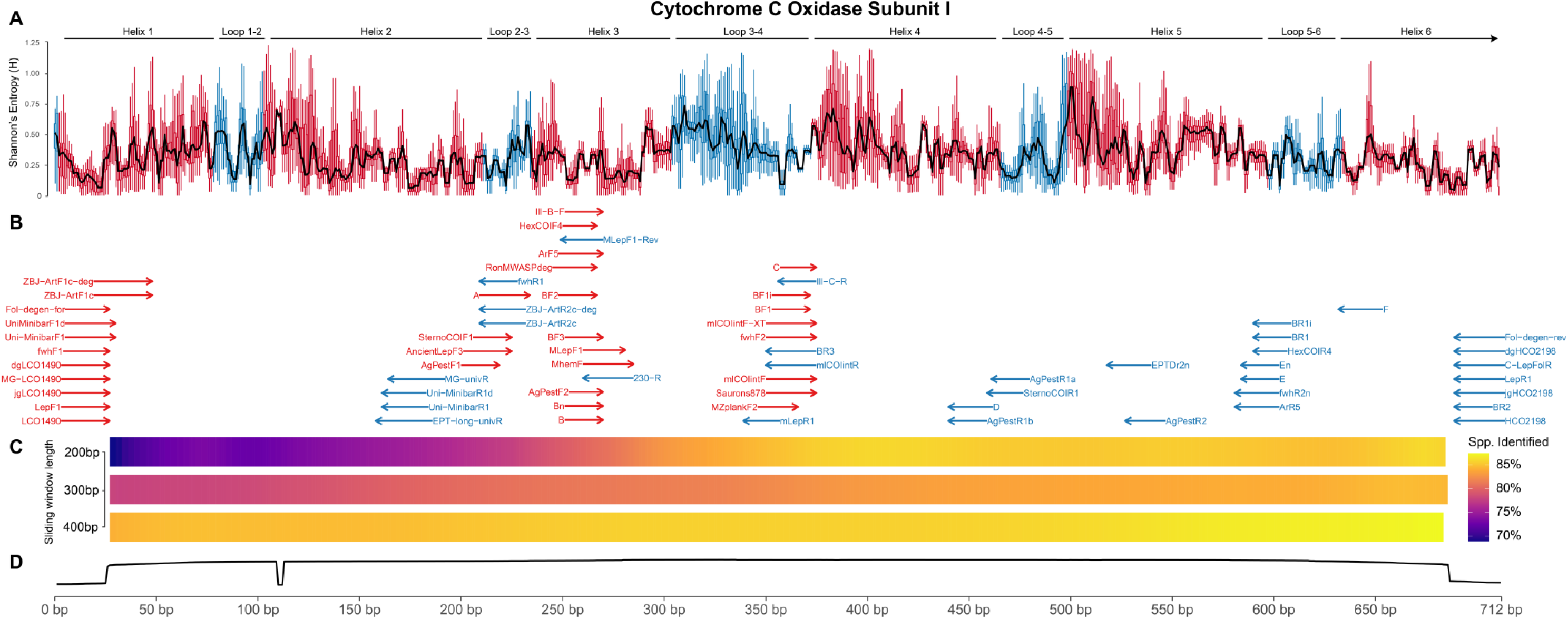
Summary of the COI barcode region. **A)** Boxplots of Shannon’s entropy per nucleotide position calculated separately for each Insect order in the database, with structural motifs annotated as per Pentinsaari et al. (2016). **B)** Binding positions of all published and novel primers evaluated in this study. **C)** Identification success for all insects within sliding windows of length 200 bp, 300 bp and 400 bp. **D)** Number of sequences at each position within the COI barcode locus.

### Diagnostic sensitivity of mini-barcodes

PCR amplification was simulated by truncating the reference sequences to the region amplified by each primer set as originally published, as well as the 1156 unique combinations of forward and reverse primers that could produce an amplicon >50 bp. To determine how the percentage identity threshold used to assign species impacted identification success, virtual amplicons were clustered at the commonly used identification threshold of 97% (Alberdi et al., 2018), as well as more stringent 98%, 99%, and 100% thresholds using *UCLUST* (Edgar, 2010). Again, each species was considered successfully identified if there were no sequences with conflicting taxonomic annotations within its respective cluster. As the relationship between identification success and amplicon length was not linear, a segmented regression model was fit separately to each distance interval using the *chngpt* R package (Fong et al., 2017). The changepoint between the two regression segments, which can be considered the minimum amplicon length at which identification success using COI mini-barcodes becomes congruent with the full-length barcode region, was determined via maximum likelihood with confidence intervals obtained from 1000 bootstrap replicates (Fong, 2019). Primer specific identification performance was obtained by averaging the residuals from the regression model for all evaluated combinations that contained that respective primer. In order to determine how the taxa that could not be identified by any barcode length were distributed across the insect phylogeny, the consenTRAIT metric (Martiny et al., 2013) was calculated for the full-length barcode at the 97% identification threshold, with each clade weighted by the number of failed identifications. This metric measures the mean phylogenetic depth of clades for which a binary trait, in this case failed identification, is present in at least 50% of its tips, with significance assessed against 1000 permutations. The phylogenetic clades with the highest number of identification failures were then annotated with their lowest common taxonomic rank.

### Primer-template mismatch

A mismatch score was calculated between each primer and every sequence that contained an appropriate binding site using *PrimerMiner* (Elbrecht & Leese, 2017a). The default penalties as per Stadhouders et al., (2010) were used to score types of mismatches, with penalty scores doubled for each contiguous mismatch and increased exponentially towards 3’ end of primer. To determine the phylogenetic scale at which primer-template mismatch is conserved, 1×10^8^ pairs of tips were randomly selected from the phylogenetic tree and binned into 100 discrete intervals of phylogenetic distance. The phylogenetic autocorrelation function (Zaneveld & Thurber, 2014), or how the value of a trait (primer mismatch score) decays with increasing phylogenetic divergence was then calculated separately for each primer using the *castor* R package (Louca & Doebeli, 2018). As primer-template mismatch was found to be phylogenetically conserved (supplementary Figure 4), phylogenetic independent contrasts (Felsenstein, 1985) was used to impute mismatch scores for species that did not have available sequence data within their respective primer binding sites (Zaneveld & Thurber, 2014). The accuracy of phylogenetic imputation is determined by the depth at which the trait is conserved (measured by the decay of the autocorrelation function), as well as the distance from the tip being imputed to the nearest clade with available data, which was quantified for each primer set using the Nearest Sequence Taxon Index (NSTI) averaged over all tips in the phylogeny (Langille et al., 2013). To determine the importance of primer degeneracy for reducing bias, a linear regression model was fit separately to the imputed and unimputed mismatch data for all forward and reverse primers.

### Ranking of primers

To obtain an overall ranking for each evaluated primer, the identification success for both the pest and entire insect dataset, as well as the inverse of the primer mismatch score were z-normalised to be on the same scale. Additionally, each forward and reverse primer was assigned a value of either −1 (poor), 0 (moderate), or 1 (good), depending on how closely their physical characteristics adhered to common primer design recommendations (Supplementary Table 2) (Abd-Elsalam, 2003; Kwok et al., 1994; Shen et al., 2010), and these were also normalised. The standardised scores from each metric were then weighted by their relative importance for overall primer performance; 1x for mismatch, 0.5x for pest insect identification, 0.5x for all insect identification, and 0.25x for each of the measured physical characteristics; fold-degeneracy, primer length, melting temperature, GC%, presence of GC clamps and longest homopolymer, then summed by primer to obtain a final ranking.

## Results

### Sequence database assembly

To assemble the reference database for primer evaluation, 4,491,128 COI sequences with taxonomy “Insecta” were retrieved from GenBank and BOLD, including 23,571 extracted from mitochondrial genomes. Of these sequences, 1,584,589 were exclusive to GenBank, 15,153 were exclusive to BOLD, and 2,891,386 shared across both repositories (Figure 1B). 2,745,595 sequences and 129,225 species could be mapped to valid binomial names within the OTT taxonomy, including 11,431 taxonomic synonyms resolved to currently accepted species names in the process. This step resulted in the largest reduction of both unique sequences and species (Figure 1B), with most unsuccessfully mapped sequences being flagged as incertae_sedis (1,329,337 sequences) or not present in the taxonomy at all (346,299 sequences). Additional reasons for removal at this stage included infraspecific taxa (22,501 sequences), the taxon being extinct (4,493 sequences) or other more minor issues (supplementary Figure 1). In contrast, many of the later curation steps only marginally reduced the number of both sequences and species (Figure 1B), with homology and pseudogene filters removing 333,765, and 16,553 sequences respectively, and comparisons of taxonomy between highly similar sequences removing 3,130 putatively misannotated sequences. A BLAST search against a local database of contaminants removed 30 Wolbachia and 56 human mitochondrial sequences and identified a further 1,667 sequences that were >98% similar to known COI pseudogenes but did not contain any characteristic stop codon or frameshift mutations. All sequences <200 bp in length were then removed, leaving 2,389,404 sequences from 110,676 distinct species remaining. When compared to globally relevant lists of invasive insects, a total of 2,625 species spanning 1,490 genera and 20 taxonomic orders were identified within the curated database (supplementary Figure 2). Each pest species was represented by a significantly higher number of sequences (mean 79.9 ± 5.60) than the average insect taxon (mean 20.2 ± 0.428) (Welch t-test: t_(2837)_ = 11.1, p < .001), however no reference data was available for 1,717 of the listed species. As the number of sequences per species was greatly skewed by a few highly sampled species (supplementary Figure 3), the database was pruned to a maximum of 5 sequences per taxa. This left a total of 315,754 sequences from 110,676 species remaining in the final curated reference database (Figure 1B).

### Optimal diagnostic subregions within the COI barcode

The final curated reference database consisted of a 712 bp alignment (Figure 2), which included the 709 bp barcode region and a 3bp insertion at position 110-112 that occurred in the order Thysanoptera as well as some Hymenopteran species. Sequence coverage was relatively even across the COI barcode, with exception of the terminal regions where the standard practice of removing priming sequences before submission to public repositories resulted in low coverage (Figure 2D). The per-site Shannon entropy within regions coding for loop structures was significantly higher than within those coding for transmembrane helices (Welch t-test: t_(12306)_ = 4.0459, p < .001), however short segments of low entropy were distributed throughout both (Figure 2A). The large majority of the 58 primers retrieved from the literature were designed around these few low-entropic segments, with many overlapping around 180 bp, 255 bp, 370 bp, and 600 bp into the alignment (Figure 2B). As none of these segments of low entropy extended for the 18-24 bp length of a typical primer, many of the evaluated primers included multiple degenerate bases to account for variable positions (Supplementary Table 1). Sliding window analyses revealed that for a 200 bp or 300 bp amplicon, those positioned towards the 3’ end of the barcode region could differentiate substantially more species than those towards the 5’ end, however differences were much less pronounced for a 400 bp amplicon (Figure 2C).

### Diagnostic sensitivity of mini-barcodes

PCR amplification was simulated for the 1156 primer combinations that could produce an amplicon >50 bp, as well as the full-length barcode region. Identification success using mini-barcodes generally increased with amplicon length, but did not follow a simple linear relationship, and instead saw a sharp initial increase up to a certain length, followed by a second more gradual slope (Figure 3). A segmented regression model applied to the whole insect dataset inferred the change point between these trends to be 257 bp at the 97% identity threshold (95% CI: 242 bp – 266 bp), 237 bp at the 98% threshold (95% CI: 218 bp – 248 bp), 146 bp at the 99% threshold (95% CI: 137 bp – 206 bp) and 125 bp when only 100% matches were considered (95% CI: 110 bp – 131 bp). On the other hand, the inferred changepoints for the pest dataset were approximately 20 bp smaller at the 97%, 98% and 99% identity thresholds, but identical to the larger dataset at the 100% threshold. In spite of this general trend, certain amplicons deviated up to 10% above or below the regression line for both datasets (Figure 3), reinforcing that appropriate placement of mini-barcodes within the COI barcode region can be just as important as amplicon length for diagnostic performance.

**Figure 3:**
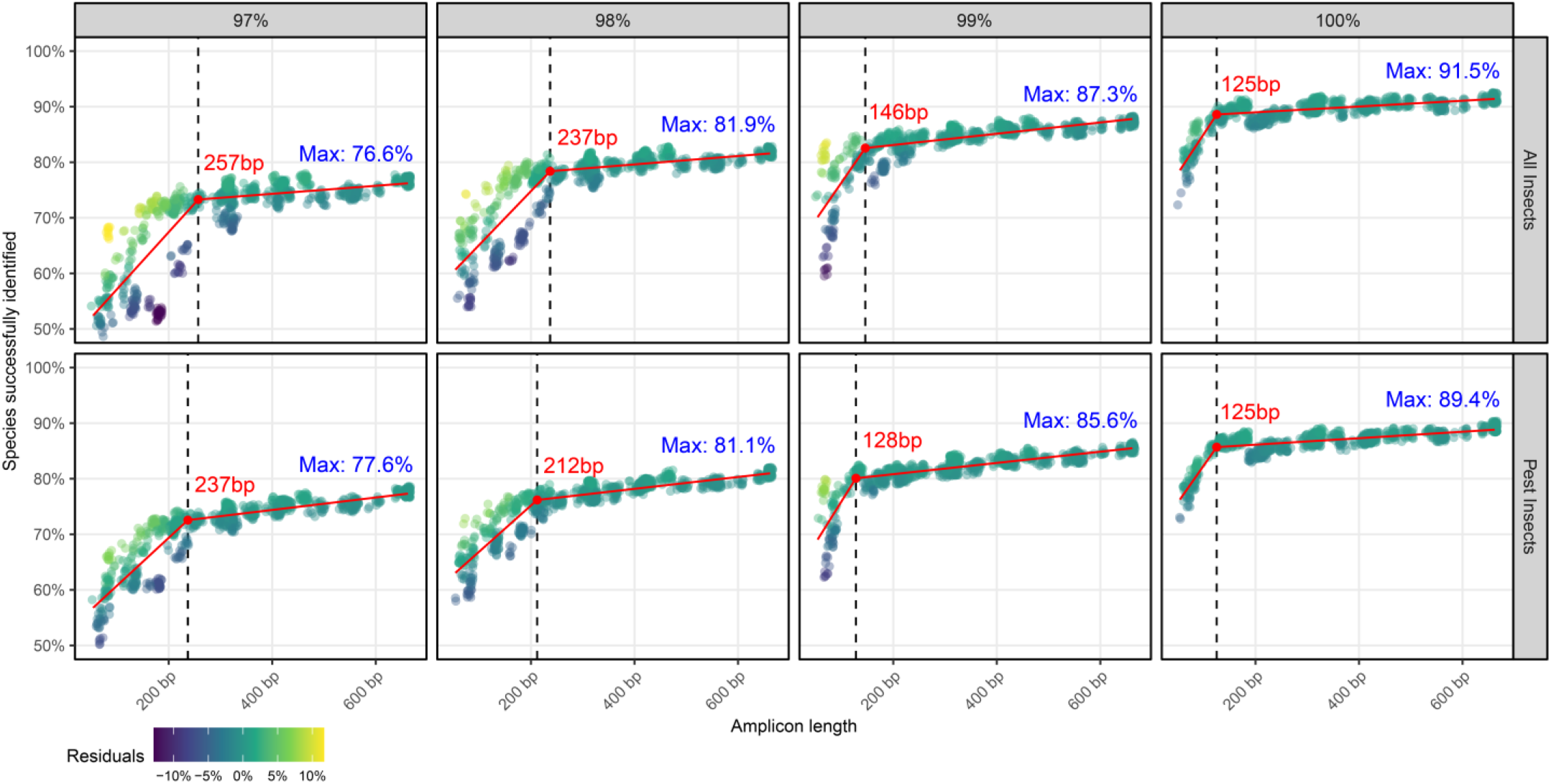
Identification success for all insects (upper) and Insect pests (lower) as a function of amplicon length for all possible combination of mini-barcode primers. Percentage identity threshold used to identify species increases from 97% (leftmost panel) to 100% (rightmost panel), with segmented regression model predictions overlaid. Each primer combination is coloured by its residual error from regression model predictions, with higher residuals indicating better than expected performance for its length.

The overall proportion of Insect species that could be successfully identified by any of the barcodes increased with the stringency of the percentage identity threshold used (Figure 3). For the complete Insect dataset, the proportion identified increased from 76.6% when using a 97% identification threshold up to 91.5% when only 100% matches were considered, and similarly increased from 77.6% to 89.4% in the pest dataset. Taxa that could not be identified even by the full-length barcode showed significant phylogenetic clustering (Figure 4A), with the mean phylogenetic depth of problem clades (those in which ≥50% of species couldn’t be differentiated from their congenerics) ranging from 0.83% at a 97% identity threshold (consenTRAIT, p < .001) to 0.21% at 100% identity threshold (p < .001). These patterns indicate that identification failure with DNA barcoding can be considered a phylogenetically conserved trait, but concentrated in smaller clades scattered throughout the tips of the phylogeny rather than being inherent to any broader lineage (Figure 4A). These problem clades mostly occurred within the order Lepidoptera, which contained the highest number of failed species identifications (Figure 4C). In particular, the families *Noctuidae, Nymphalidae*, and *Tortricidae* each contained several large problematic clades (Figure 4B). Other non-Lepidopteran taxa that posed problems for DNA barcode based identification included Hoverflies (Diptera: *Syriphidae*), sawflies (Hymentopeta: *Empria*) and the hemipteran families *Aphididae* and *Lachnidae* (Figure 4B), however to a substantially lesser degree than Lepidoptera (Figure 4C).

**Figure 4:**
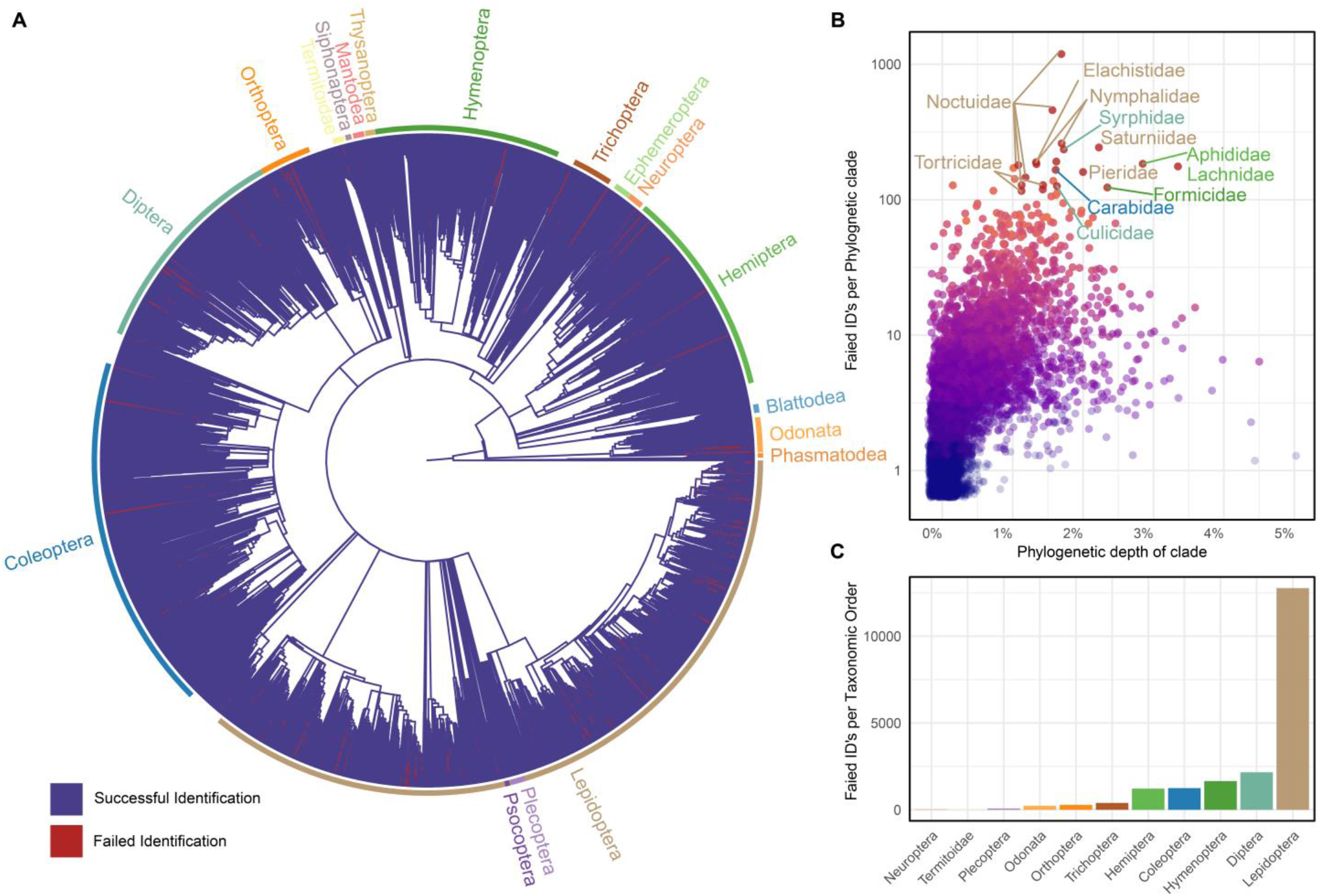
**A)** Phylogenetic tree of all insect genera contained within the curated reference database with major orders annotated. Clades are highlighted in red where ≥50% of species could not be successfully differentiated from their congenerics by the full-length barcode a 97% identity threshold. **B)** Phylogenetic depth of clades where ≥50% of specimens could not be identified, with those containing the highest number of failed identifications annotated. **C)** Number of failed identifications per taxonomic order with the full-length barcode region at the 97% identity threshold.

### Predicted bias for metabarcoding primers

Primer-template mismatch scores were calculated between the 68 primers and all taxa that had reference sequences available for their respective binding sites, and scores for missing species phylogenetically imputed. Mismatch was found to be phylogenetically conserved across the majority of primers, with the autocorrelation function decaying moderately with phylogenetic divergence to reach a correlation of 0.5 around a phylogenetic distance of 5%, and falling to zero at a distance of 7-10% (supplementary Figure 4). This indicates that missing data imputation would be reliable for those taxa within 5% phylogenetic divergence from a sequenced clade, and close to random for species greater than 7% diverged. Almost all primers had a mean phylogenetic distance to their nearest sequenced taxon (NSTI) between 0.05% and 0.35%, well below this value. The exception was those primers situated at each terminus of the COI barcode where available sequence data was sparser (Figure 2), leaving the NSTI between 4% and 6% (Figure 5D). Following imputation, the forward primers with the lowest mean mismatch across all insect taxa were: C, BF1i, ARF5, and the reverse primers ArR5, E, EN and BR1 (Figure 5B). Nevertheless, these primers still showed significant mismatch to certain clades within the phylogenetic tree, most notably the families *Diaspididae, Pseudococcidae, Philopteridae, Phlaeothripidae, Apidae, Gyrinidae*, Leiodidae, *Hesperiidae* and the Coleopteran genus *Exapion* (Figure 5C). At a higher level, the orders Hymenoptera and Hemiptera showed substantially more mismatching taxa then any of the other major insect groups (Figure 5A, B, C). Primer-template mismatch was found to be significantly related to primer degeneracy for both the imputed (p < .001, R^2^ = .187) and unimputed datasets (linear regression, p < .001, R^2^ = .152), yet those primers with the highest degeneracy were not necessarily the best performers (Figure 5D). The diminishing returns of adding degeneracy was particularly apparent for the ZBJ-ArtF1c-deg, mtCOIF-XT and MZPlankF2 forward primers and the reverse primer D, which despite extremely high degeneracy still showed moderate mismatch across the insect phylogeny (Figure 5D). The underperformance of highly degenerate primers such as ZBJ-ArtF1c-deg remained when only the unimputed data was viewed (Supplementary Figure 5), suggesting that these results were not confounded by difficulties imputing higher levels of missing data.

**Figure 5:**
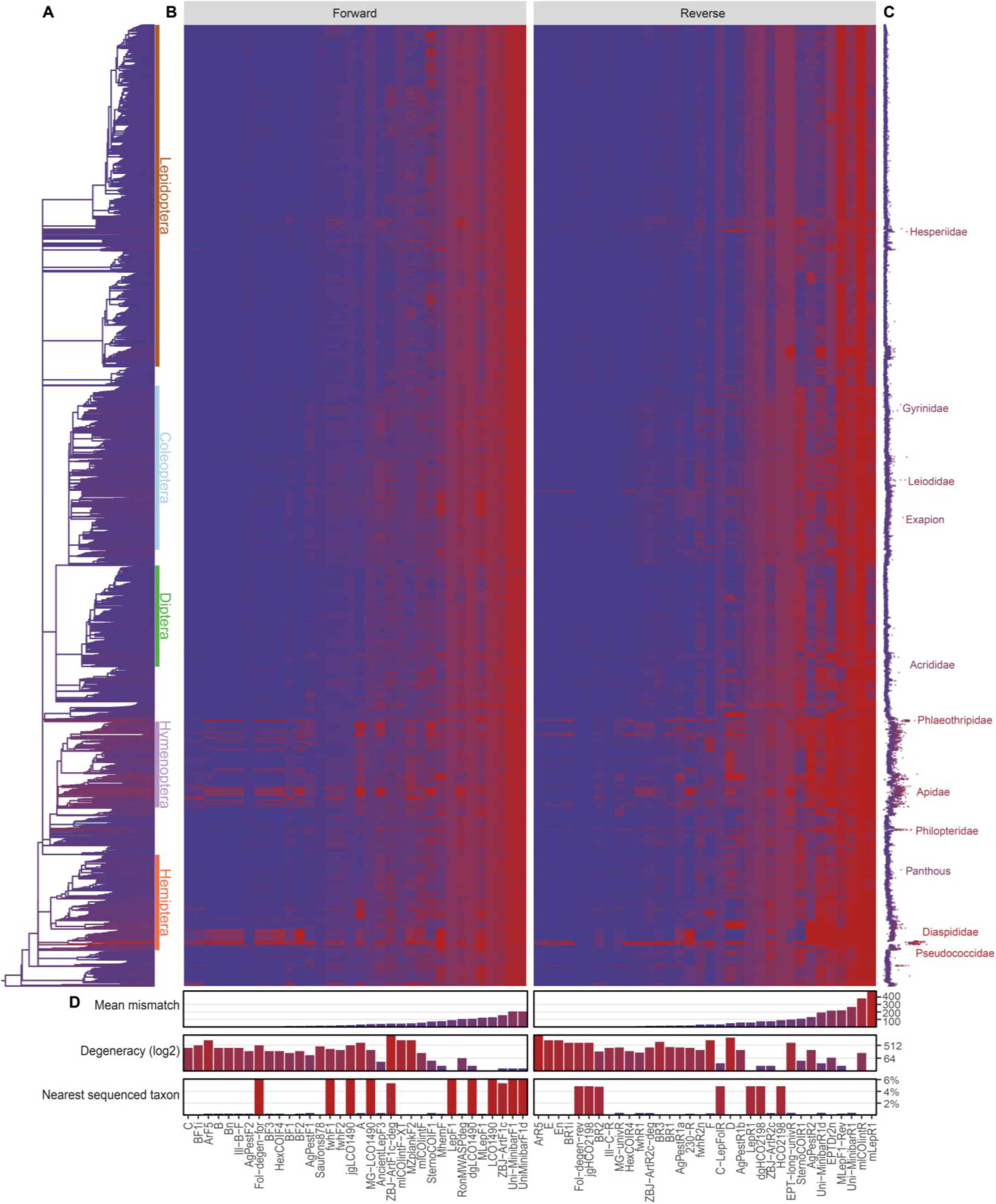
**A)** Phylogenetic tree of all insect genera contained within the curated sequence database, coloured by their mean primer-template mismatch across all evaluated primers. **B)** Primer-template mismatch for each evaluated sequence and primer, summarised by insect genus. **C)** Dotplot of mean mismatch per genus, with highly mismatched clades indicated. **D)** Summary of results by primer, from top to bottom; mean primer mismatch across all sequences, fold degeneracy on log2 scale, mean phylogenetic distance from each imputed tip to its nearest sequenced taxon.

### Final primer rankings

Many of the evaluated forward primers performed well across all metrics, with C, BF1, fwhF2, BF1i, SauronS878 and MlCOIintF all ranking highly (Figure 6). Nevertheless, there was no perfect forward primer, with many of those showing higher diagnostic sensitivity also having excessive mismatches, or physical characteristics outside of recommended design guidelines (Figure 6). In contrast, there was substantially more variability in the performance of reverse primers, with primers such as D, AgPestR1a and AgPestR1b showing exceptional diagnostic sensitivity, but having either too much degeneracy or a melting temperature well below the recommended guidelines. On the other hand, the reverse primers Ill-C-R, fwhR2n, and BR2 showed slightly less sensitivity, but had no critical issues for any of the measured physical characteristics (Figure 6). To comply with restrictions of 2 × 150 bp sequencing, the primer combinations fwhF2-fwhR2n, BF1-BR1, or SauronS878-fwhR2n present the best overall options, amplifying a ~250 bp subregion of COI that contains the most diagnostic nucleotides (Figure 2A, C), while showing little mismatch across all insects and physical characteristics within recommended guidelines. The novel AgPestF1-AgPestR1b primers designed in this study would also provide an appropriate 260 bp amplicon for 2 × 150 bp sequencing, however further laboratory evaluation would be required to ensure their lower than recommended melting temperatures does not introduce non-specific amplification. On the other hand, a much greater range of primer combinations are appropriate for sequencing technologies that can deliver read lengths of 2 × 300 bp. In particular, those which amplify a subregion from 250 bp into the full-length barcode, along to either the low entropy region around 600 bp, or onwards the 3’ terminus will capture the most diagnostic nucleotides (Figure 2C). Published primer combinations that amplify these subregions and performed well in the evaluated metrics include HexCOIF4-HexCOIR4, BF2-BR2 and BF3-BR2, but many of the alternative forward or reverse primers that overlap the same positions would also prove suitable (Figure 2B). Nevertheless, once the amplicon has reached approximately 400 bp there is minimal difference in species discrimination across the COI barcode (Figure 2C), and therefore primer combinations that amplify from the 5’ end of the barcode onwards would likely also perform adequately for 2 × 300 bp sequencing.

**Figure 6:**
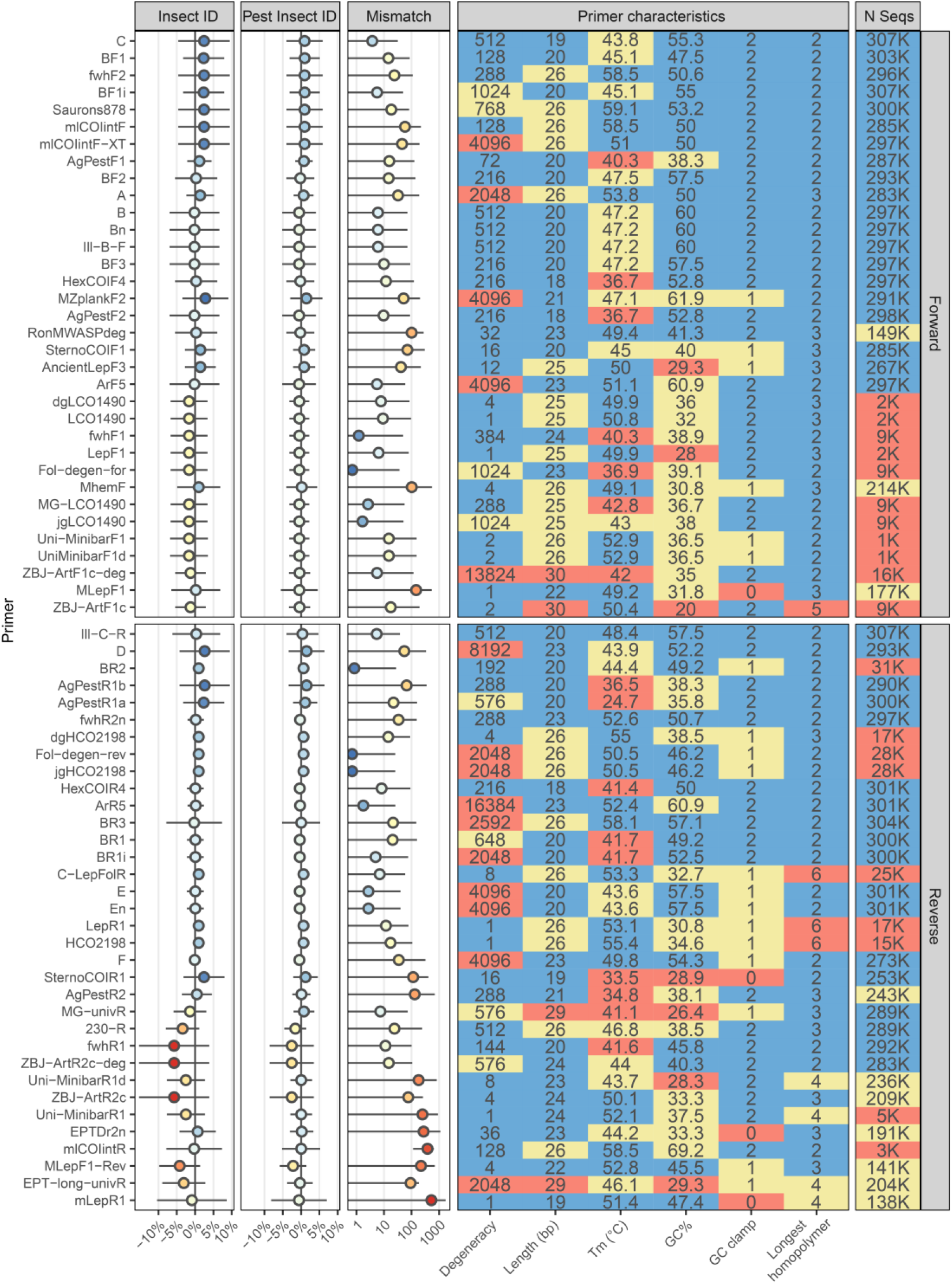
Summary of primer performance across all metrics measured within this study, with primers ranked from best overall performance (top) to worst (bottom). Metrics from left to right: Average performance of primer compared to regression model predictions on all insects and pest insects, mean mismatch score for each primer, and primer characteristics. Error bars represent 2 standard deviations.

## Discussion

High sensitivity across a broad taxonomic scope is a defining feature of DNA metabarcoding assays, facilitating their use as a universal diagnostic assay to rapidly screen mixed samples for a range of target pests or pathogens. Despite several studies applying metabarcoding to certain pest taxa (Batovska et al., 2018, 2020; Bowser et al., 2019; Young et al., 2021), the diagnostic performance of the required mini-barcodes has not until now been systematically evaluated across the broader diversity of invasive insect species. Using a curated reference database covering 110,676 insect species, including 2,625 species registered on global invasive species lists, we here demonstrate that mini-barcodes can achieve comparable resolution to the full-length COI barcode region already widely accepted within insect diagnostic protocols. Our findings are largely in agreement with previous investigations showing congruence between mini-barcodes and the full-length barcode (Hajibabaei, Smith, et al., 2006; Meusnier et al., 2008), as well as morphospecies (Yeo et al., 2020). However, our study expands these predictions to a more than five-fold larger sample of insect taxa, with an additional focus on invasive insect pests.

While 97% identity is considered the default threshold for delineating taxonomic units from DNA barcodes (Alberdi et al., 2018; Porter & Hajibabaei, 2020), our analyses demonstrate that a more stringent 98% or 99% identification threshold not only increases the number of insects that can successfully be differentiated, but also reduces the amplicon length required to do so. This is particularly notable for implementing metabarcoding on production scale HTS platforms such as the Illumina NovaSeq, which offer the lowest cost per sample but require much shorter amplicons due to their typical read lengths of only 2 × 150 bp (Piper et al., 2019). While use of more stringent identification thresholds has been constrained by the common practice of clustering metabarcoding reads to resolve sequencing errors (Porter & Hajibabaei, 2020), recent denoising algorithms provide single nucleotide resolution that can be leveraged for more accurate and reproducible taxonomic assignment (Callahan et al., 2017; Porter & Hajibabaei, 2020). Nevertheless, the metric of identification success used within our study (proportion of clusters containing only a single species) does not consider the actual availability of reference sequences to match unknown taxa against, and false negatives may be introduced when using these more stringent thresholds if intraspecific diversity isn’t sufficiently represented in the reference database. While this will not be an issue for most invasive insect pests due to their general overrepresentation in public sequence repositories, some taxa were represented by only single sequences, and a further 1,717 had no publicly available COI data whatsoever. Moreover, the 110,676 insect species represented in our curated database barely accounts for 10% of described insect diversity (Stork, 2018), highlighting the considerable efforts still required to increase the taxonomic coverage of reference databases before metabarcoding assays can operate in a truly universal manner.

While many of the evaluated mini-barcodes performed comparably to the full-length barcode region, 10.6% of insect pests, and 8.5% of all insect species could not be differentiated at all, even when solely considering 100% matches. While this may at first glance seem high, it is largely consistent with previous research that suggests between 12.3% and 26.5% of described Arthropod species are non-monophyletic for the COI barcode region (Funk & Omland, 2003; Mutanen et al., 2016; Ross, 2014). In our study, these misidentified taxa were phylogenetically clustered in “problem clades” towards the tips of the phylogeny, which predominantly occurred within the order Lepidoptera. Issues of DNA barcode failure for Lepidoptera and other speciose taxonomic groups has long been noted (Meier et al., 2006; Wiemers & Fiedler, 2007), however only recently has it been appreciated how much of this can be attributed to underlying misidentifications, databasing errors, or flawed taxonomic delimitation (Locatelli et al., 2020; Mutanen et al., 2016). While our study has followed current best practices in reference database curation, we notably did not go to the extra length of verifying the original source for the taxonomic identities applied to each sequence, a task which would prove insurmountable for a dataset of this size. While computational curation presents an extremely scalable approach for resolving annotation errors and contamination within public datasets, the quality of any identification ultimately depends on the quality of the systematics and taxonomy that originally delimited and described the species (Clarke & Schutze, 2014). Therefore, additional taxonomic consideration using more comprehensive genomic (Leaché et al., 2014) or integrative methods (Padial et al., 2010) may be required to determine if the problem clades identified in our study are actually due to insufficient resolution within the COI barcode, or rather, over-splitting in the underlying taxonomy (Mutanen et al., 2016).

The 11,431 defunct or synonymous species names identified and corrected in our study clearly demonstrates that taxonomic synonyms remain one of the largest and seldom discussed issues within public sequence repositories (Leray et al., 2019). While taxonomic names must be free to change to reflect revised species concepts, this becomes a problem when historically defunct species names are retained in reference databases, propagating errors through later studies and the management decisions made from them (Clarke et al., 2019). For insect metabarcoding, issues arising from taxonomic synonyms will become most apparent as hierarchical taxonomic classifiers already widely adopted by microbiome researchers become more prevalent (Porter et al., 2014; Porter & Hajibabaei, 2018). These methods require a query sequence to reach a certain bootstrap support to be assigned to lower ranks in the taxonomy, but conflicts in taxonomic annotation between genetically similar reference sequences could result in a failure to reach the required confidence, and thus false negative detections. Therefore, we recommend that resolving taxonomic synonyms to their currently accepted name become a default step in all metabarcoding database curation efforts, alongside the more common practices of removing non-homologous sequences, contaminants, and misannotated taxonomy (Kozlov et al., 2016; Richardson et al., 2020). While some curation of taxonomic synonyms already occurs within both GenBank and BOLD (Schoch et al., 2020), determining the currently valid taxonomic name from a diverse and constantly evolving primary literature is by no means a trivial task (Schoch et al., 2017). Continued investment into digital infrastructure for distribution of taxonomic information will therefore prove critical for ensuring identifications obtained through metabarcoding remain robust to the inevitable future description and renaming of taxa (Miralles et al., 2020).

In addition to its ability to differentiate target species, the amount of PCR amplification bias towards or against certain taxonomic groups plays an important role in the selection of primers for metabarcoding studies. While the lack of evolutionarily conserved primer binding sites led to early studies questioning the suitability of COI for metabarcoding (Deagle et al., 2014), our analyses demonstrate that incorporating a moderate 216 to 512-fold degeneracy into primers (4-5 degenerate bases) can adequately resolve primer-template mismatch across the large majority of insect taxa, with diminishing returns beyond this. While not explicitly evaluated in this study, previous research has shown that primers with over 2000-fold degeneracy are likely to cause undesired amplification of non-target taxa (Collins et al., 2019), a particular problem for samples with low DNA concentrations (Leese et al., 2021; Macher et al., 2018). With this in mind, we advise against the use of extremely degenerate primers such as ArR5, ZBJ-ArtF1c-deg, or D for insect metabarcoding, in favour of other primers that overlap similar regions of COI and show similar performance despite substantially less degeneracy. Of the taxonomic groups that still showed a high level of mismatch despite inclusion of degenerate bases, the most concerning for an invasive species surveillance programme would be the Armoured Scales (Hemiptera: *Diaspididae*), Mealybugs (Hemiptera: *Pseudococcidae*), and the thrips family *Phlaeothripidae*, for which 120, 90, and 28 species respectively were registered on global invasive species lists. While *Apidae* also showed substantial mismatch across many primers, this was only towards non-pest taxa within the family, and most well-designed primers matched the 26 invasive *Bombus* and *Apis* species well. In all these cases, researchers and diagnosticians should be aware that primer-template mismatches could cause false negatives when these taxa are at a low relative abundance within mixed trap samples. Therefore, the sequencing effort applied to each sample may need be adjusted in proportion to both the overall biomass and expected taxonomic composition of the communities under study, if known in advance.

Despite this study being a purely in-silico evaluation and the many limitations that this entails (Alberdi et al., 2018; Corse et al., 2019; Zhang et al., 2020), it represents a comprehensive first step in applying big data principles to inform development of HTS based diagnostics for invasive insect pests. As well as providing a starting point for diagnosticians and researchers selecting mini-barcode primers for insect identification, our results offer a degree of confidence to managers and regulators grappling with the consequences of non-targeted HTS assays and how to appropriately respond the incidental detection of regulated species (Darling et al., 2020). Whilst we still recommend additional laboratory validation be conducted on high-priority targets before adoption in active surveillance, promisingly, many of the outstanding primers highlighted in our rankings have also shown similarly high performance in a recent in-vitro evaluation on a diverse insect mock community (Elbrecht et al., 2019). In summary, appropriately chosen COI mini-barcode primers perform effectively across the majority of pest and non-pest insects, opening for the adoption of universal metabarcoding assays within diagnostic laboratories. While the computational curation pipeline presented here can resolve many issues inherent to public reference sequence data, regardless of whether the diagnostic tool is a microscope or a HTS assay, the accuracy of results will ultimately depend on the underlying quality and completeness of the taxonomy for the target groups.

## Acknowledgements

The authors would like to thank Francesco Martoni, Conrad Trollip and Brendan Rodoni for valuable discussions on metabarcoding and taxonomy. Financial support for this study was provided by Agriculture Victoria’s Improved Market Access for Horticulture programme (CMI105584) and the Horticulture Innovation Australia iMapPESTS project (ST16010) through funding from the Australian Government Department of Agriculture as part of its Rural R&D for Profit program and Grains Research and Development Corporation. A.M.P. is supported by Australian Government Research Training Program Scholarship.

## Author contributions

A.M.P. conceptualised the study, performed all analyses, and drafted the manuscript with input and supervision from N.O.I.C., J.P.C., and M.J.B. All authors read and approved the final version of the manuscript.

## Availability of data and materials

Functions and a tutorial for curating COI reference databases are provided in the ‘taxreturn’ R package, available on GitHub: https://github.com/alex-piper/taxreturn All code required to reproduce the statistical analyses and figures presented in this paper are contained within the following GitHub repository: https://github.com/alexpiper/primer_evaluation

