## Supplementary information for "Computational Evaluation of DNA Metabarcoding for Universal Diagnostics of Invasive Insect Pests"

bioRχiv

### Supplementary material

| Primer | Strand | Sequence | Citation |
|---|---|---|---|
| **SternoCOIF1** | F | ATTGGWGGWTTYGGAAAYTG | Batovska et al. (2021) |
| **SternoCOIR1** | R | ATRAARTTRATWGCTCCTA | Batovska et al. (2021) |
| **Saurons878** | F | GGDRCWGGWTGAACWGTWTAYCCNCC | Rennstam Rubbmark et al. (2018) |
| **AgPestF1** | F | ATYATWATTGGDGGDTTYGG | This Study |
| **AgPestF2** | F | HGAYATRGCHTTYCCHCG | This Study |
| **HexCOIF4** | F | HCCHGAYATRGCHTTYCC | Marquina et al. (2019) |
| **HexCOIR4** | R | TATDGTRATDGCHCCNGC | Marquina et al. (2019) |
| **mLepR1** | R | CCTGTTCCAGCTCCATTTT | Hebert et al. (2004) |
| **AgPestR1a** | R | GTRATRAARTTDAYWGMHCC | This Study |
| **AgPestR1b** | R | ARAATWGADGADAYWCCWGC | This Study |
| **AgPestR2** | R | RACWGMTCAVAYAAATARDGG | This Study |
| **LCO1490** | F | GGTCAACAAATCATAAAGATATTGG | Folmer et al. (1994) |
| **HCO2198** | R | TAAACTTCAGGGTGACCAAAAAATCA | Folmer et al. (1994) |
| **Uni-MinibarR1** | R | GAAAATCATAATGAAGGCATGAGC | Meusnier et al. (2008) |
| **Uni-MinibarR1d** | R | AAAATTATAATAAARGCRTGRGC | Jordaens et al. (2013) |
| **Uni-MinibarF1** | F | TCCACTAATCACAARGATATTGGTAC | Meusnier et al. (2008) |
| **UniMinibarF1d** | F | TCCACTAATCACAARGATATTGGTAC | Jordaens et al. (2013) |
| **ZBJ-ArtF1c** | F | AGATATTGGAACWTTATATTTTATTTTTGG | Zeale et al. (2011) |
| **ZBJ-ArtF1c-deg** | F | RGAYATYGGWACHYTWTAYTTYHTHTTYGG | Elbrecht et al. (2019) |
| **ZBJ-ArtR2c** | R | WACTAATCAATTWCCAAATCCTCC | Zeale et al. (2011) |
| **ZBJ-ArtR2c-deg** | R | WAYTARTCARTTWCCRAAHCCHCC | Elbrecht et al. (2019) |
| **mlCOIintF** | F | GGWACWGGWTGAACWGTWTAYCCYCC | Leray et al. (2013) |
| **mlCOIintR** | R | GGRGGRTASACSGTTCASCCSGTSCC | Leray et al. (2013) |
| **BR3** | R | GGDGGRTANACWGTYCAHCCDGTHCC | Elbrecht et al. (2019) |
| **LepF1** | F | ATTCAACCAATCATAAAGATATTGG | Hebert et al. (2004) |
| **EPT-long-univR** | R | AARAAAATYATAAYAAANGCGTGNANNGT | Hajibabaei et al. (2011) |
| **MLepF1-Rev** | R | CGTGGAAAWGCTATATCWGGTG | Brandon-Mong et al. (2015) |
| **Ill-C-R** | R | GGNGGRTANACNGTTCANCC | Shokralla et al. (2015) |
| **Ill-B-F** | F | CCNGAYATRGCNTTYCCNCG | Shokralla et al. (2015) |
| **BF1** | F | ACWGGWTGRACWGTNTAYCC | Elbrecht & Leese (2017b) |
| **BF1i** | F | ACNGGNTGRACNGTNTAYCC | Elbrecht et al. (2019) |
| **BF2** | F | GCHCCHGAYATRGCHTTYCC | Elbrecht & Leese (2017b) |
| **BF3** | F | CCHGAYATRGCHTTYCCHCG | Elbrecht et al. (2019) |
| **BR1** | R | ARYATDGTRATDGCHCCDGC | Elbrecht & Leese (2017b) |
| **BR1i** | R | ARYATNGTRATNGCNCCNGC | Elbrecht et al. (2019) |

| | | | |
|---|---|---|---|
| **BR2** | R | TCDGGRTGNCCRAARAAYCA | Elbrecht & Leese (2017b) |
| **ArF5** | F | GCNCCNGAYATRKCNTTYCCNCG | Gibson et al. (2014) |
| **ArR5** | R | GTRATNGCNCCNGCNARNACNGG | Gibson et al. (2014) |
| **jgLCO1490** | F | TNTCNACNAAYCAYAARGAYATTGG | Geller et al. (2013) |
| **jgHCO2198** | R | TANACYTCNGGRTGNCCRAARAAYCA | Geller et al. (2013) |
| **MZplankF2** | F | RGYNGGNACRGGNTGRACNGT | Elbrecht et al. (2019) |
| **LepR1** | R | TAAACTTCTGGATGTCCAAAAAATCA | Hebert et al. (2004) |
| **C-LepFolR** | R | TAAACTTCWGGRTGWCCAAAAAATCA | Hernández-Triana et al. (2014) |
| **AncientLepF3** | F | TTATAATTGGDGGWTTTGGWAATTG | Prosser et al. (2016) |
| **A** | F | GGNGGNTTTGGNAATTGAYTNGTNCC | Hajibabaei et al. (2012) |
| **D** | R | CCTARNATNGANGARAYNCCNGC | Hajibabaei et al. (2012) |
| **B** | F | CCNGAYATRGCNTTYCCNCG | Hajibabaei et al. (2012) |
| **Bn** | F | CCNGAYATRGCNTTYCCNCG | Elbrecht et al. (2019) |
| **E** | R | GTRATNGCNCCNGCNARNAC | Hajibabaei et al. (2012) |
| **En** | R | GTRATNGCNCCNGCNARNAC | (Elbrecht et al. (2019) |
| **C** | F | GNTGAACNGTNTAYCCNCC | Hajibabaei et al. (2012) |
| **F** | R | CCNGCNGGRTCNAARAANGANGT | Hajibabaei et al. (2012) |
| **fwhF1** | F | YTCHACWAAYCAYAARGAYATYGG | Vamos et al. (2017) |
| **fwhR1** | R | ARTCARTTWCCRAAHCCHCC | Vamos et al. (2017) |
| **fwhF2** | F | GGDACWGGWTGAACWGTWTAYCCHCC | Vamos et al. (2017) |
| **fwhR2n** | R | GTRATWGCHCCDGCTARWACWGG | Vamos et al. (2017) |
| **MG-LCO1490** | F | ATTCHACDAAYCAYAARGAYATYGG | Galan et al. (2018) |
| **MG-univR** | R | ACTATAAARAARATYATDAYRAADGCRTG | Galan et al. (2018) |
| **230-R** | R | CTTATRTTRTTTATNCGNGGRAANGC | Gibson et al. (2015) |
| **MhemF** | F | GCATTYCCACGAATAAATAAYATAAG | Park et al. (2011) |
| **dgHCO2198** | R | TAAACTTCAGGGTGACCAAARAAYCA | Meyer (2003) |
| **dgLCO1490** | F | GGTCAACAAATCATAAAGAYATYGG | Meyer (2003) |
| **Fol-degen-for** | F | TCNACNAAYCAYAARRAYATYGG | D. W. Yu et al. (2012) |
| **Fol-degen-rev** | R | TANACYTCNGGRTGNCCRAARAAYCA | D. W. Yu et al. (2012) |
| **MLepF1** | F | GCTTTCCCACGAATAAATAATA | Hajibabaei, Janzen et al. (2006) |
| **RonMWASPdeg** | F | GGWTCWCCWGATATAKCWTTTCC | Clare et al. (2019) |
| **mlCOIintF-XT** | F | GGWACWRGWTGRACWNTNTAYCCYCC | Wangensteen et al. (2018) |
| **EPTDr2n** | R | CAAACAAATARDGGTATTCGDTY | Leese et al. (2021) |

**Supplementary Table 1:** Published and novel primers evaluated in this study

| **Criterion** | **Good (1)** | **Moderate (0)** | **Bad (-1)** |
|---|---|---|---|
| **GC%** | 40%-60% | 30%-40% or 60%-70% | <30% or >70% |
| **Degeneracy** | 0-517 fold | 517-1026 fold | >1026 fold |
| **GC clamp (last 2 bases of 5' end)** | 2 G or C bases | 1 G or C base | No G or C bases |
| **Primer length** | 18-24 bp | 16-18 bp or 24-26 bp | <16 bp or >25 bp |
| **Longest homopolymer** | ≤2 bp | ≤4 bp | >4 bp |
| **Melting temperature** | 48-62 °C | 43-48 °C or 63-67 °C | <43 °C or >67 °C |

**Supplementary Table 2:** Criteria used to rank primer characteristics

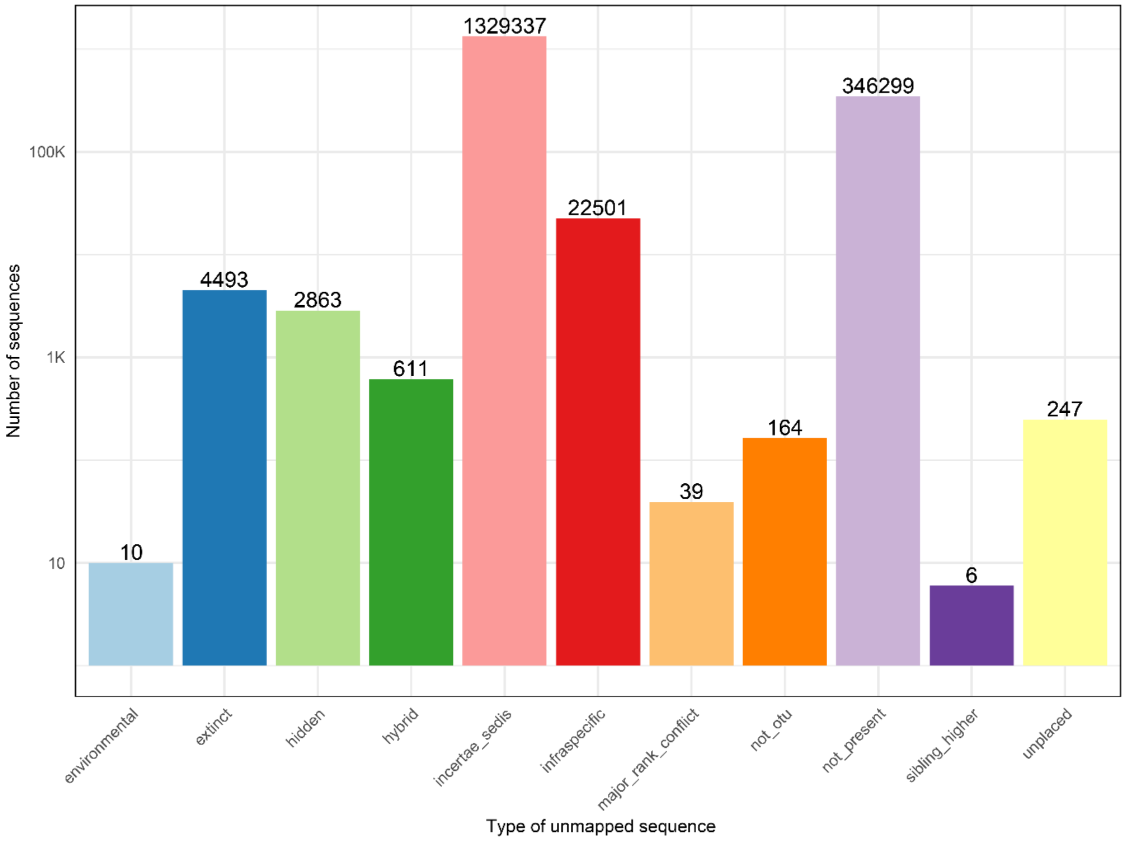


**Supplementary Figure 1:** Categories of sequences that could not be successfully mapped into the Open Tree of Life taxonomy.

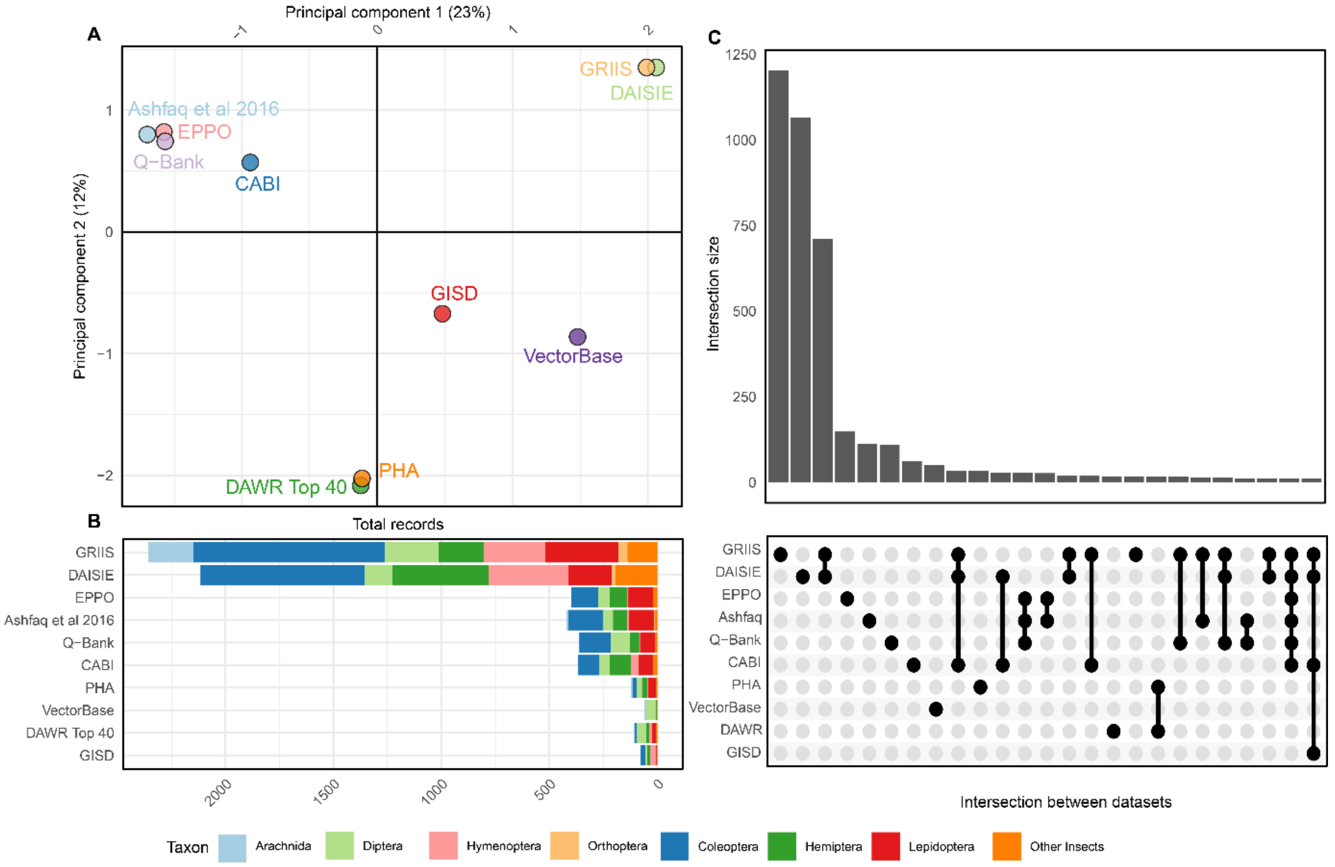


**Supplementary Figure 2:** Summary of public pest and invasive insect datasets used to assemble the pest list for primer evaluation. A) Principal component analysis of species overlap (Jaccard distance) between pest lists. B) Total records for each dataset source, coloured by taxonomic order or class. C) 25 largest intersections of species names between invasive or pest species datasets.

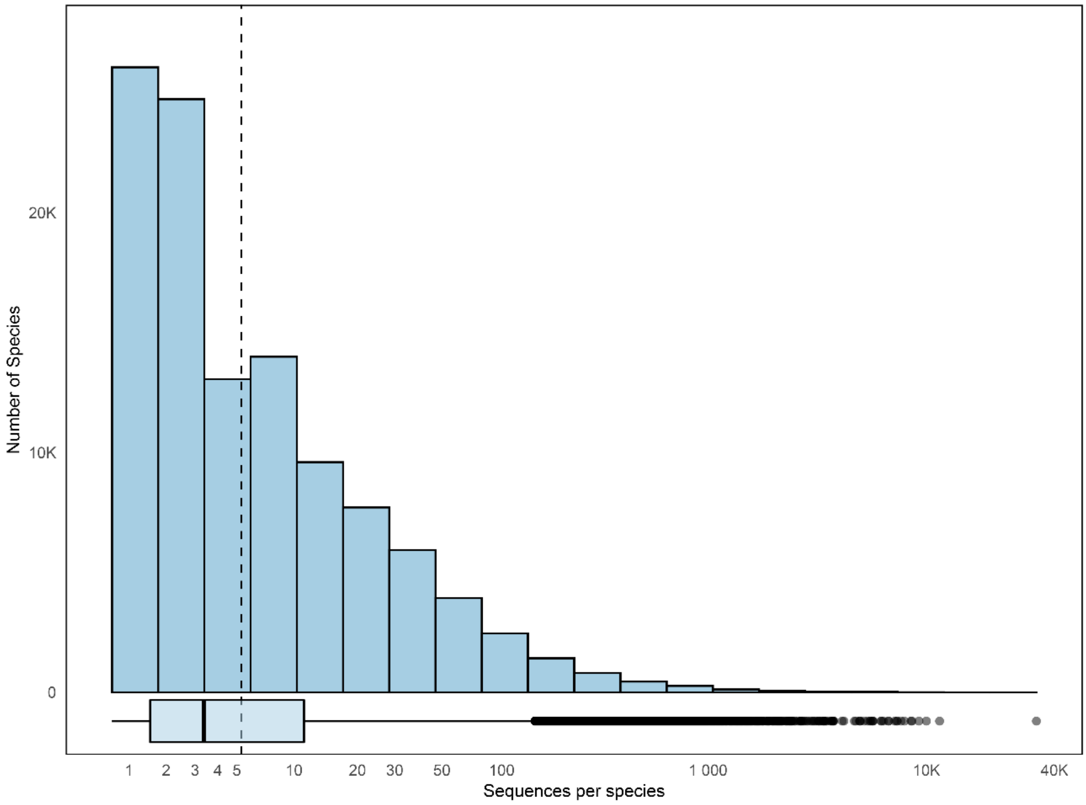


**Supplementary Figure 3:** Number of sequences per species in the reference database before pruning. Vertical line indicates the maximum of 5 sequences per species the final database was pruned to.

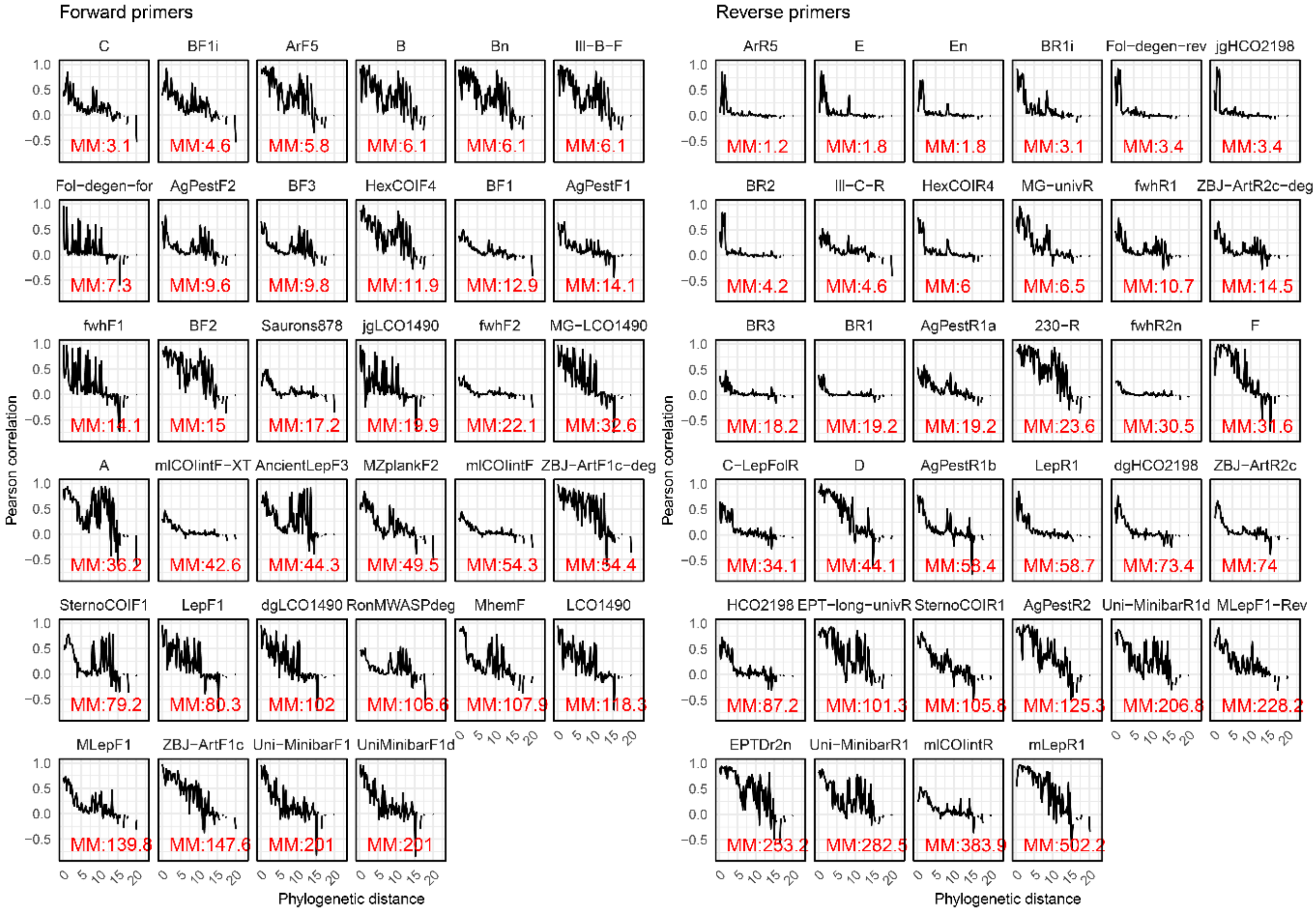


**Supplementary Figure 4:** Phylogenetic autocorrelation function of primer mismatch for each forward and reverse primer. Annotations refer to mean mismatch between primer and all insect species.

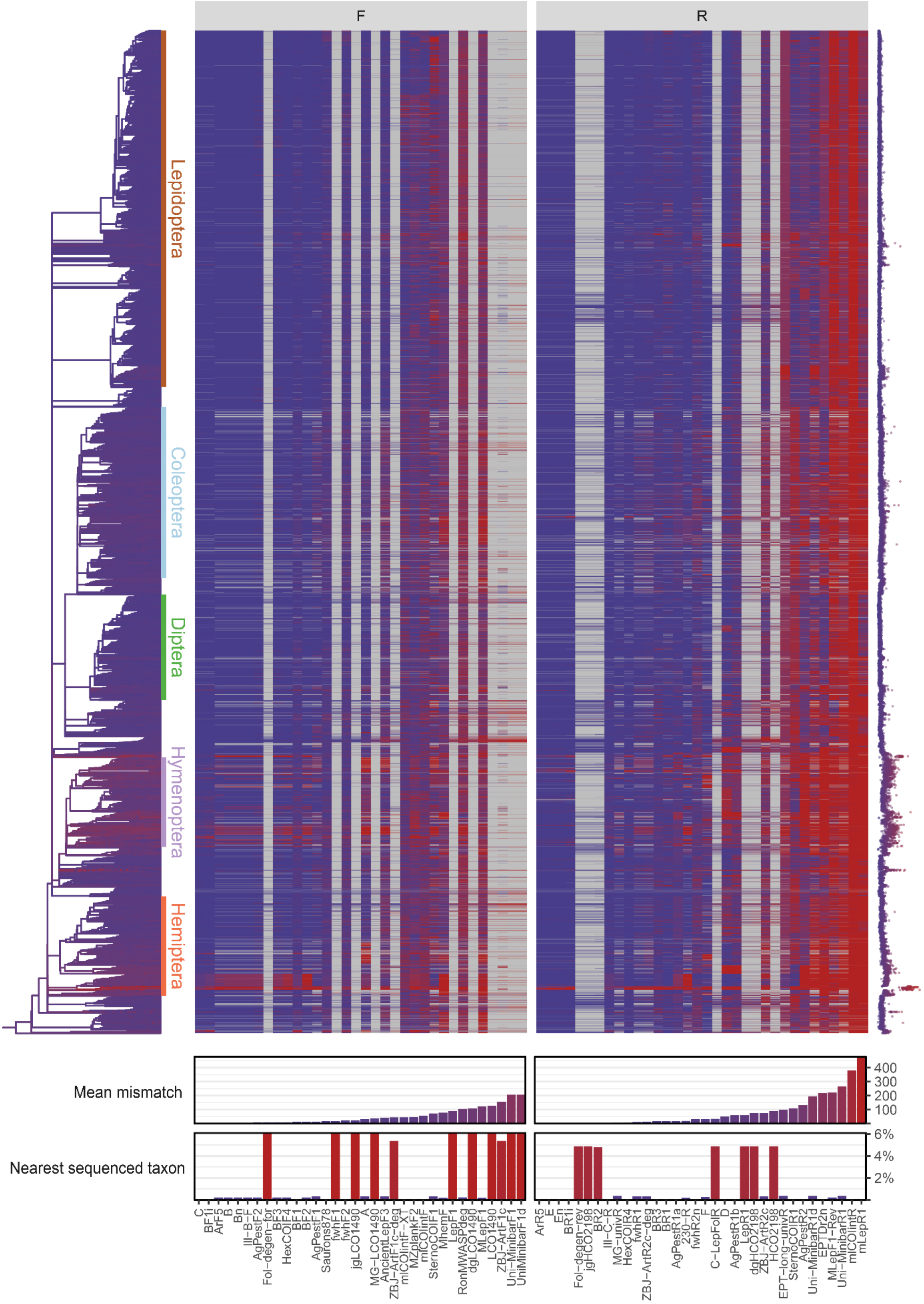


**Supplementary Figure 5:** Alternative to main figure 5 without imputation of missing data

**Supplementary Note 1:** Sources for invasive or pest species records used to assemble global list of pest Arthropods.

- EPPO global database: https://gd.eppo.int/
- US APHIS: https://www.aphis.usda.gov/aphis/home/
- QBank: https://qbank.eppo.int/arthropods/organisms
- Global invasive species database: http://www.iucngisd.org/gisd/search.php
- Global register of introduced or invasive species: http://www.griis.org/
- VectorBase: https://www.vectorbase.org/organisms
- DAWR top 40: http://www.agriculture.gov.au/pests-diseases-weeds/plant
- PHA National biosecurity status report: http://www.planthealthaustralia.com.au/national-programs/national-plant-biosecurity-status-report/
- Ashfaq & Herbert 2016: DNA barcodes for bio-surveillance: regulated and economically important arthropod plant pests
- CABI: https://t.co/LGjlFoOazd
- http://www.europe-aliens.org

**Supplementary Note 2:** All sequences that were mapped to nodes within the Open Tree of Life that were annotated with these flags that indicate uncertain placement were removed during sequence filtering.

- incertae_sedis
- major_rank_conflict
- infraspecific
- unplaced
- environmental
- inconsistent
- extinct
- hidden
- hybrid
- not_otu
- viral
- barren

**Supplementary Note 3:** All sequences with taxonomic annotations containing these words that indicate insufficient identification were removed during sequence filtering.

- sp.
- spp.
- aff.
- nr.
- bv.
